# A reverse-transcription/RNase H based protocol for depletion of mosquito ribosomal RNA facilitates viral intrahost evolution analysis, transcriptomics and pathogen discovery

**DOI:** 10.1101/453910

**Authors:** Joseph R. Fauver, Shamima Akter, Aldo Ivan Ortega Morales, William C. Black, Americo D. Rodriguez, Mark D. Stenglein, Gregory D. Ebel, James Weger-Lucarelli

## Abstract

Studies aimed at identifying novel viral sequences or assessing intrahost viral variation require sufficient sequencing coverage to assemble contigs and make accurate variant calling at low frequencies. Many samples come from host tissues where ribosomal RNA represents more than 90% of total RNA preparations, making unbiased sequencing of viral samples inefficient and highly expensive, as many reads will be wasted on cellular RNAs. In the presence of this amount of ribosomal RNA, it is difficult to achieve sufficient sequencing depth to perform analyses such as variant calling, haplotype prediction, virus population analyses, virus discovery or transcriptomic profiling. Many methods for depleting unwanted RNA or enriching RNA of interest have been devised, including poly-A selection, RNase H based specific depletion, duplex-specific nuclease treatment and hybrid capture selection, among others. Although these methods can be efficient, they either cannot be used for some viruses (i.e. non-polyadenylated viruses), have been optimized for use in a single species, or have the potential to introduce bias. In this study, we describe a novel approach that uses an RNaseH possessing reverse transcriptase coupled with selective probes for ribosomal RNA designed to work broadly for three medically relevant mosquito genera; *Aedes*, *Anopheles,* and *Culex.* We demonstrate significant depletion of rRNA using multiple assessment techniques from a variety of sample types, including whole mosquitoes and mosquito midgut contents from FTA cards. To demonstrate the utility of our approach, we describe novel insect-specific virus genomes from numerous species of field collected mosquitoes that underwent rRNA depletion, thereby facilitating their detection. The protocol is straightforward, relatively low-cost and requires only common laboratory reagents and the design of several small oligonucleotides specific to the species of interest. This approach can be adapted for use with other organisms with relative ease, thus potentially aiding virus population genetics analyses, virus discovery and transcriptomic profiling in both laboratory and field samples.

## Introduction

The past several decades have witnessed the emergence and expansion of viruses with increasing frequency (Jones et al., 2008). Several examples are H1N1 Influenza in 2009 (Otte et al., 2015), Chikungunya in 2006 (Tsetsarkin et al., 2007), Zika in 2013-14 (Aubry et al., 2017), West Nile in 1999 (Moudy et al., 2007), MERS (Forni et al., 2015) and others. Most, if not all of the emerging viruses that pose the greatest threat to human and animal health are RNA viruses. In fact, all 7 of the pathogens identified by the World Health Organization (WHO) in the 2018 annual review of the blueprint list of priority diseases as requiring urgent or serious research were RNA viruses, with the other being unknown pathogens (WHO, 2018). Due to the importance of RNA viruses, it is critical to be able to detect, identify and analyze these pathogens’ genomes using novel high-throughput sequ byencing methods. However, total RNA preparations from complex lab and field samples typically contain extremely high levels of ribosomal RNA (rRNA), sometimes 80-90% of the total amount (Eun, 1996). Sequencing data mapping to rRNA is typically removed bioinformatically and therefore represents an economic waste and reduces the number of samples that can be tested in a given sequencing run. To increase the number of reads mapping to sequences of interest, several methods have been employed to either enrich; hybrid-capture (Metsky et al., 2017), amplicon (Metsky et al., 2017; Moratorio et al., 2017), SPIA amplification (Grubaugh et al., 2016) or remove unwanted rRNA sequences; ribosomal RNA depletion most notably (Adiconis et al., 2013; Matranga et al., 2016). Enriching for sequences of interest is highly effective but can only target for specific sequences, making it difficult to identify novel, divergent viruses. Current ribosomal depletion methods are typically cost-prohibitive or are only effective for human and mice, making depletion in non-model systems highly inefficient.

Here we describe a novel method of ribosomal depletion that utilizes reverse transcription (RT) to specifically target sequences for depletion using the RNA degradation activity of RNase H. During RT, rRNA is converted to cDNA using specific DNA probes which can then be degraded using RNase H. Because this method utilizes small probes to recognize the target sequences for depletion, it is possible to design universal probes that bind to highly divergent species, making depletion of diverse organisms representing several genera possible with the same probes. Additionally, since the probes are effectively reverse primers that are typically used for RT, they are easy to design, cheap and can be quickly designed for any target sequence from any species or genus of interest for which the rRNA sequence is available. Our studies show that using this method we can selectively remove rRNA from mosquitoes from multiple genera which results in increased relevant data recovered from next-generation sequencing. Furthermore, we apply this method to field-caught mosquitoes and show that we are able to detect multiple novel virus genomes from a highly multiplexed set of samples on a relatively low-output Illumina MiSeq run. Collectively, this work describes an effective method for rRNA depletion that is straight forward, relatively low-cost and highly effective at increasing usable data from high-throughput sequencing experiments.

## Materials and Methods

### Cells, Viruses, Mosquitoes and Sample Collection

West Nile virus (strain NY99) was generated from an infectious clone as previously described in BHK-21 cells (Shi et al., 2002). Laboratory colonies of *Culex quinquefasciatus, Aedes aegypti* and *Anopheles gambiae* were used for mosquito infections. Mosquitoes were maintained at 26-27°C and 70-80% relative humidity with a 16:8 L:D photoperiod. Water and 10% sucrose were provided *ad libitum*. For preliminary studies, pools of whole mosquitoes (n=10) were collected and homogenized in Trizol solution.

For xenosurveillance studies, groups of *An. gambiae* were exposed to an infectious bloodmeal containing 10^7^ PFU of WNV NY99. The next day, midguts from mosquitoes containing a residual bloodmeal were collected by spreading the midgut contents onto CloneSaver FTA cards (GE Healthcare), and immediately 25μL of RNAlater (ThermoFisher) was added in order to facilitate diffusion of blood into the FTA card and stabilize the nucleic acid. The samples placed on the FTA cards were then punched out and nucleic acid was eluted by incubation in RNA rapid extraction solution (ThermoFisher) for 18 hours.

### Ribosomal Depletion

A more detailed protocol is included describing the depletion protocol in Supplemental file 1. Nucleic acids were extracted using either Trizol solution or the Mag-Bind Viral DNA/RNA kit (Omega Bio-tek, USA) and eluted into 50μL of water. The samples were then treated with TURBO DNase (ThermoFisher) and purified using RNAClean XP beads (Beckman Coulter). For reverse transcription, the RNA was mixed with oligos specific for rRNA (sequences listed in Supplemental Table 1) and dNTPs and then heat denatured at 95C for 2 minutes followed by slow cooling to 50C at 0.1C/s. For initial experiments, we tested a panel of reverse transcriptases (RTs), including *Tth* DNA polymerase (in the presence of Mn2+, Promega), Superscript III (SSIII, ThermoFisher), Superscript IV (SSIV, ThermoFisher), Avian myeloblastosis vi-rus (AMV, NEB) and Moloney Murine Leukemia Virus (MMLV, NEB). For all RTs, we used the optimal conditions as described by the manufacturer. For all further experiments, AMV RT was then added and incubated at 50C for 2 hours. RNase H (NEB) was then added to destroy the RNA present in the RNA:cDNA hybrid. The samples were then digested with DNase I (NEB) to remove the cDNA and residual oligos. The RNA was then purified using RNAClean XP beads at a 1.8x ratio.

### RNA Analysis and qRT-PCR

Input and rRNA depleted RNA were analyzed using a 2100 Bioanalyzer (Agilent) per manufacturer’s protocols with the total RNA pico kit. The RNA traces were analyzed using Agilent 2100 expert software. Quantitative Reverse-transcriptase PCR (qRT-PCR) was performed using the iTaq universal probes supermix (Biorad) according to the manufacturer. qRT-PCR was performed with the following primers; 18S Forward - AGAGGACTACCATGGTTGCAAC, 18S Reverse - CCTGCTGCCTTCCTTGGATG, 18S Probe - CCGGAGAGGGAGCCTGAGAAATGGC, 28S Forward - AGGTGCGGAGTTTGACTGG, 28S Reverse - TCCTTATGCTCAGCGTGTGG, 28S Probe - AGGTGTCCAAAGGTCAGCTCAGTGTGG, WNV Forward - TCAGCGATCTCTCCACCAAAG, WNV Reverse - GGGTCAGCACGTTTGTCATTG, WNV Probe - TGCCCGACCATGGGAGAAGCTC (Lanciotti et al., 2000). The number of genome copies was generated by fitting the Ct values to a standard curve of RNA specific to each of the primer sets.

### Library Preparation and Data Analysis

Libraries for Illumina sequencing were prepared from both input RNA and samples that were depleted using probes specific to rRNA or in the absence of probes. The libraries were prepared using equal concentrations of RNA as input by using the NEBNext Ultra RNA library prep kit (NEB) and then were sequenced on an Illumina MiSeq using 150 cycles. For data analysis, libraries were first demultiplexed using bcl2fastq (Illumina). Reads were then trimmed for both adapters and quality using BBDuk software (part of the BBMap suite, https://sourceforge.net/projects/bbmap/). PCR duplicates were then removed using clumpify (also part of the BBMap suite) and unique reads were mapped to reference genomes using bowtie2 (Langmead and Salzberg, 2012). We then used MultiQC to quantify the percentage of reads that mapped to each reference. These percentages were graphed using GraphPad Prism version 7. To assess intrahost variation, unique reads were mapped to the Bolahun virus reference sequence using BBMap and then variants were called using LoFreq (Wilm et al., 2012). Only variants present at greater than 5% were used for analysis.

### Mosquito collections

Adult mosquitoes were collected from multiple localities in Chiapas, Mexico over the course of three weeks in August, 2016 using CDC gravid traps (John W. Hock Company), CDC Miniature light traps (BioQuip Products) and insectazookas (BioQuip Products). Mosquitoes were euthanized using triethylamine and sorted into pools of up to 25 individuals by species, sex, and collection location (Supplemental Table 2). Mosquitoes were identified to species using morphological keys (Darsie and Ward, n.d.). For groups of mosquitoes that could not be identified, multiple individuals of each group were point mounted and preserved for later identification by local experts at the Instituto Nacional de Salud Pública facilities in Mexico. Pools of mosquitoes were preserved in RNALater (Ambion) and shipped to Colorado State University (CSU).

### Processing of field collected mosquitoes

Prior to homogenization and nucleic acid extraction, mosquito pools were centrifuged and RNA later was removed. Pools were then processed as described above. All field collected mosquito pools were subjected to rRNA depletion using the same probe mixture as the laboratory experiments. Following rRNA depletion, RNA from pools was prepared for NGS using Nextera XT following manufacturer’s instructions (Illumina). Each library was dual-indexed with a unique barcode to facilitate multiplexing using the Kapa Library Amplification Kit for Illumina (Kapa BioSystems). Libraries were then quantified using the NEBNext Library Quantification Kit for Illumina (New England Biolabs) and pooled together by equal volumes. All libraries were sequenced together on a single Illumina MiSeq run using a 300 cycle (2×150) MiSeq v3 kit.

### Identification and characterization of viral sequences

Virus contigs were identified using a previously described pipeline (Cross et al., 2018; Fauver et al., 2018) (found online at https://github.com/stenglein-lab/taxonomy_pipeline). No host filtering was conducted prior to the generation of contigs, as the majority of genera sequenced to do not have a reference genome. Amino acid similarity to other virus or virus-like sequences was determined using NCBI Blastx tool against the nr database (Altschul et al., 1990) (Supplemental Table 3). Virus contigs greater than 500 b.p. were sorted into high-level clades according to Shi et al. (Shi et al., 2016). Contigs from the same species of mosquito aligning to similar viral clades were binned together in Geneious v11.0.4 and assessed for open-reading frames (ORFS) using the Find ORFs tool (Kearse et al., 2012). Following translation of complete ORFs, amino acid sequences were queried against the Conserved Domain Database v3.16 using HHpred (Zimmermann et al., 2018). Predicted domains with an e-value > 1e-5 were used for annotation. The Luteo Sobemo virus from *Ae. aegypti* was predicted to have multiple segments based on 1) homology to the most similar virus currently described, Hubei mosquito virus (Shi et al., 2016), 2) the identification of two contigs with complete ORFs, 3) similar depth of coverage across viral segments, and 4) the co-occurence of each segment in the same libraries. All putative virus genomes were described entirely using computational methods and virus isolation was not attempted.

Phylogenetic trees were created for coding complete virus genomes. The RNA dependent RNA polymerase (RDRP) gene (Luteo-Sobemo, Levi-Narna) or the whole genome (Negevirus) was used as input for blastp, and all hits with an e-value > 1e-5 were downloaded in .fasta format from NCBI. CD-Hit - c 0.90 was used to rid dataset of similar viral RDRPs sequences (Li and Godzik, 2006). Amino acid sequences were aligned using MAFFT v7.308 -auto (Katoh and Standley, 2013). Gaps and poorly aligned sequences in the multiple alignment were removed using trimAl under default settings (Capella-Gutiérrez et al., 2009). The resulting alignments were used as input to generate phylogenetic trees using PHYML with the LG substitution model and 100 bootstraps (Guindon et al., 2010). In addition, genomic sense was inferred based on placement in phylogeny.

To calculate depth of coverage, a custom database was created by species containing all viral contigs generated in this study in addition to the 45s rDNA sequence assembled from *Ae. aegypti.* Reads from each mosquito species were competitively aligned to this database using Bowtie2 under default settings (Langmead and Salzberg, 2012). The resulting SAM file was converted into BAM format, and depth of coverage at each nucleotide position was calculated using SAMtools -depth (Li et al., 2009). As well, the percentage of total reads to viruses and rRNA sequences was calculated from this database. Novel Narna-Levi virus sequences were aligned as described above, and pairwise nucleotide identity was calculated in Geneious.

### Data availability

All sequencing data has been deposited to the SRA database under BioProject SUB4694537. Novel virus genomes have been submitted to Genbank and are pending accession number assignment.

## Results

### Reverse-transcriptase (RT) mediated ribosomal RNA (rRNA) depletion is effective for mosquitoes from three medically relevant genera

The workflow for our proposed ribosomal depletion method is outlined in Figure 1. Briefly, DNase treated RNA was reverse-transcribed using DNA probes that are in the reverse-complement orientation to the sequences for mosquito sequences for the 18S, 28S and 5.8S cellular rRNA and the 12S and 16S mitochondrial rRNA sequences. In order to design probes that work against the majority of mosquitoes, we aligned sequences from several mosquito genera obtained from the SILVA rRNA database project (Quast et al., 2013). The probes were designed specifically to regions of high sequence homology and to have a melting temperature around 65℃, thereby giving them high specificity while maintaining binding to all genera. The probe sequences are presented in Supplemental Table 1 and a schematic showing the probes aligned to the *Aedes albopictus* 45S rRNA sequence is presented in Supplemental Figure 1. For the depletion, the RNA was heat denatured in the presence of the probes and slow cooled to favor specific binding of the probes to the RNA. cDNA was then synthesized using AMV reverse transcriptase (RT). It was determined that AMV RT was superior to other RTs tested in depleting 18S and 28S from *An. gambiae* mosquitoes (Fig. 2A-B). While MMLV RT was also able to significantly reduce rRNA, it also depleted WNV RNA, while AMV did not, suggesting that the depletion was highly specific (Fig. 2C). AMV and SSIII RT were the only RTs tested that significantly reduced the amount of 18S and 28S rRNA while maintaining the same amount of WNV RNA (p<0.0001 for 18S and 28S and p=0.9990 for WNV RNA all when comparing with and without probes and by One-Way ANOVA with Tukey’s correction). We continued with AMV RT because the reduction in rRNA was more dramatic and because it is less expensive than SSIII. Following AMV RT, we treated samples with RNase H and finally DNase I to degrade the RNA in the DNA:RNA hybrid and any DNA present, respectively. Using qRT-PCR, we saw a significant reduction in 18S and 28S rRNA from *An. gambiae* only when the RT and RNase/DNase steps were included and not when any steps were omitted, suggesting that the reverse transcription and RNaseH/DNase I treatment are all required for specific depletion (Fig. 2D-E, all p<0.0001 by One-Way ANOVA with Tukey’s correction as compared to the depleted group). The reduction from the DNase treated RNA to the samples not treated with RT or depletion probes is likely due to the removal of small fragments during RNAClean bead purification.

**Figure 1:**
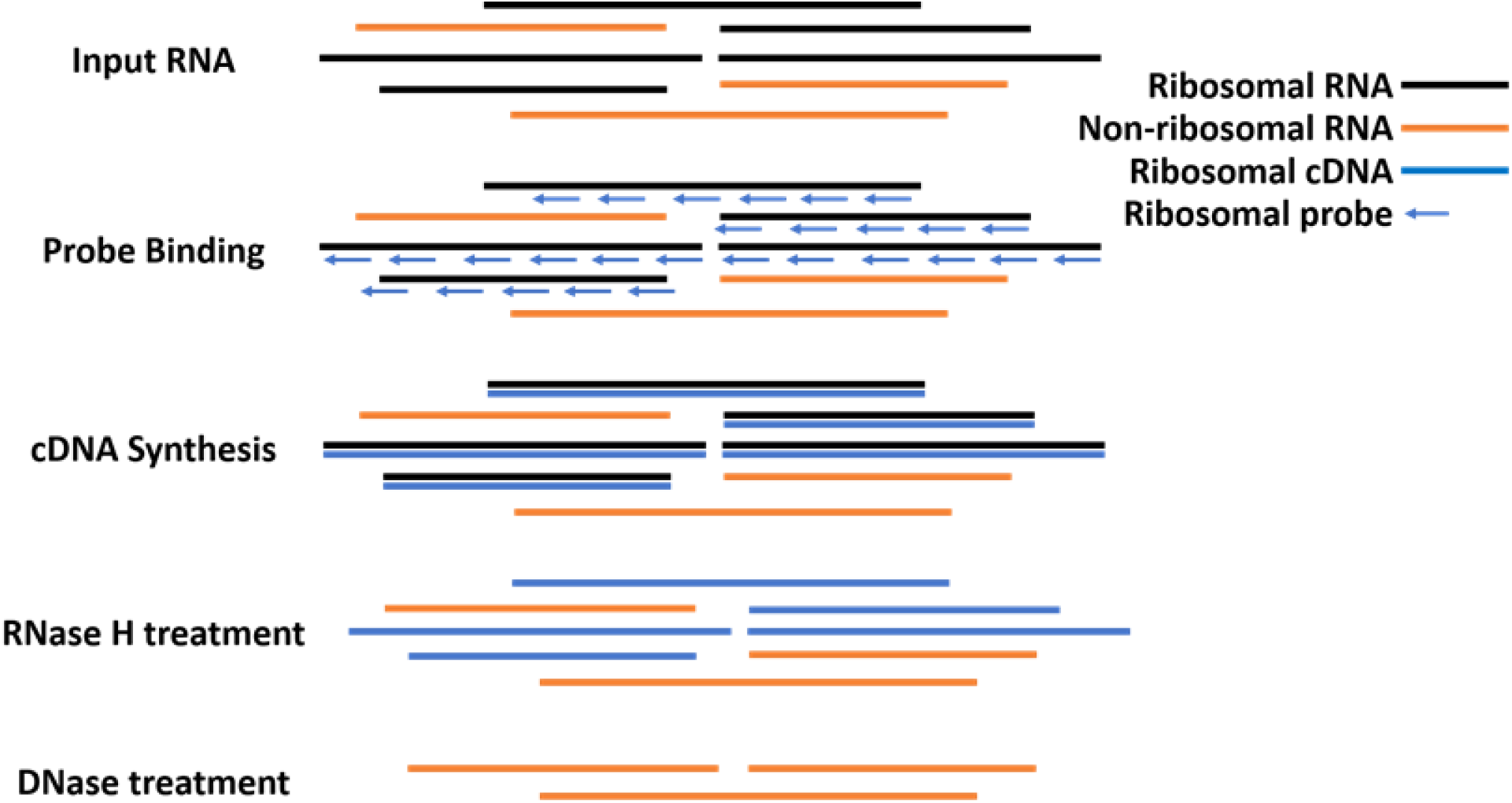
**Workflow for Reverse-Transcriptase Mediated Ribosomal Depletion from Total RNA.** To perform ribosomal RNA (rRNA) depletion, total RNA is first extracted, DNAse treated and subsequently purified with RNAClean XP Beads (Agencourt). DNA-free RNA is then bound to oligonucleotide probes designed to bind to rRNA from mosquito species in *Aedes, Culex* and *Anopheles* genera that are in the reverse complement orientation to both the long and short ribosomal subunit and 12s and 16s mitochondrial rRNA. The RNA with bound oligos is then subjected to reverse transcription using Avian Myeloblastosis Vi-rus (AMV) Reverse Transcriptase (NEB). RNA that is reverse transcribed to cDNA is then digested using RNAse H, which selectively destroys RNA in a RNA:DNA hybrid. Remaining DNA is then digested using DNAse I (NEB), leaving mostly non-ribosomal RNA which is then used for library preparation.

**Figure 2:**
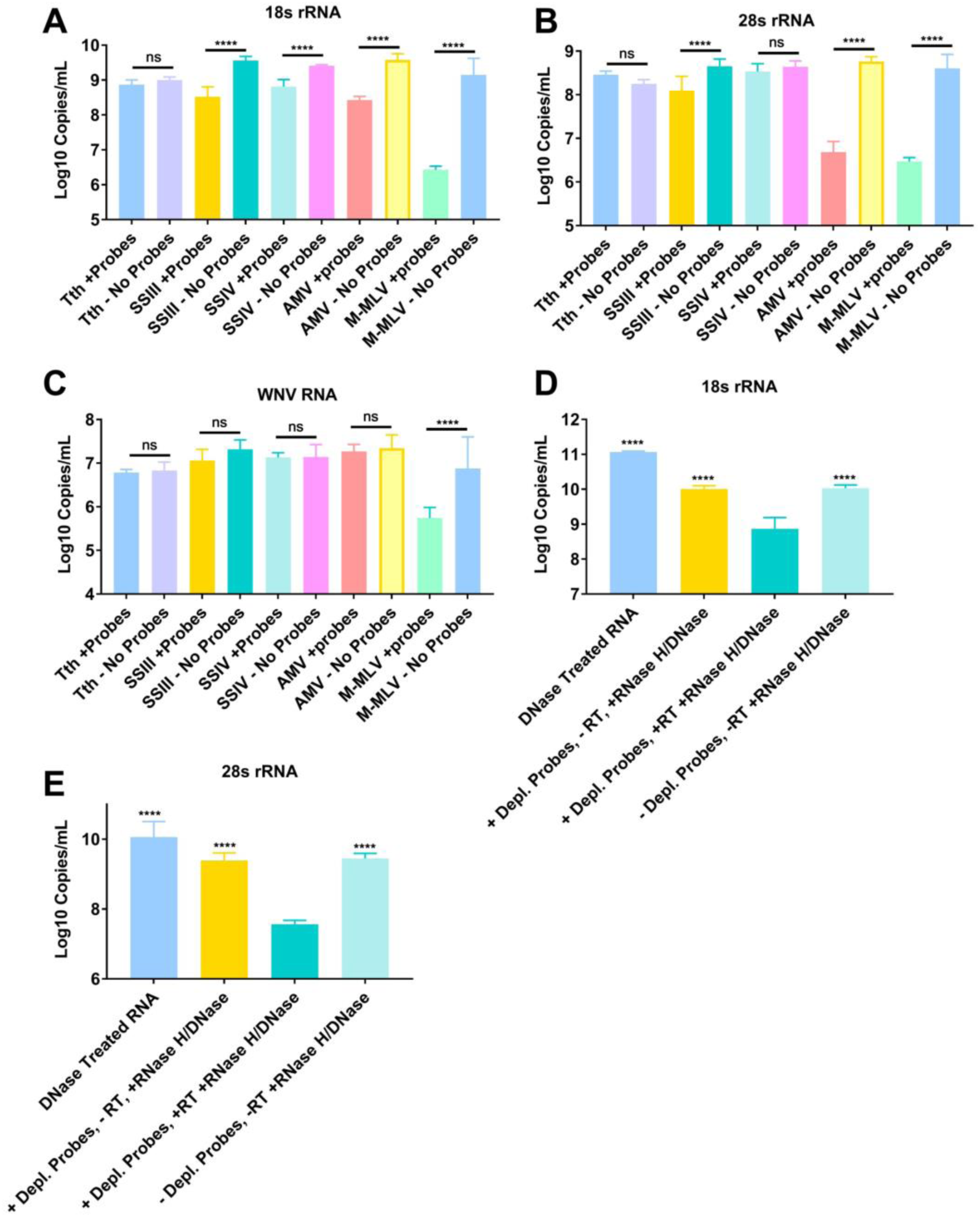
**Reverse Transcriptase mediated ribosomal RNA (rRNA) depletion is most effective with AMV RT and requires all steps to be effective**. Nucleic acids were eluted from FTA cards with midgut contents of An. gambiae that had been exposed to a bloodmeal containing West Nile virus (WNV) placed on them. RNA and DNA was then extracted to obtain total nucleic acid. The nucleic acid was then treated with DNase I and then purified to obtain total RNA. This RNA was then subjected to cDNA synthesis with a panel of reverse transcriptases in the presence (+probes) or absence (- probes) of DNA probes specific to rRNA. The RTs tested were Tth DNA polymerase, Superscript III (SSIII), Superscript IV (SSIV), AMV and MMLV. All of the samples were then treated with RNase H and then DNase I to remove the RNA present in an RNA:DNA hybrid and cDNA, respectively. The samples were then purified and subjected to qRT-PCR with primer probe combinations specific for 18S rRNA (A), 28S rRNA (B) or WNV (C). Further tests were performed exclusively with AMV RT. Panels D and E show the results of qRT-PCR for samples that underwent the process of depletion but omitting some step or reagent. 18S (D) and 28S (E) rRNA was quantified in the input RNA, RNA with no RT added, RNA with no depletion probes added and RNA treated with RT with depletion probes. All statistical tests were performed by One-Way ANOVA with Tukey’s test for multiple comparisons. **** Indicates p-value <0.0001.

We next sought to determine if the ribosomal depletion protocol was effective for mosquito species from three distinct medically relevant genera; *Anopheles, Aedes* and *Culex.* Total RNA was extracted from pools (n=10) of whole mosquitoes and the RNA was depleted as previously described, with the exception that additional probes were added to the mixture that targeted undepleted rRNA sequences identified in preliminary NGS analysis (data not shown). We then subjected the input RNA, depleted RNA (RT - with probes) and RNA that went through the depletion process without probes (RT - no probes) to qRT-PCR analysis. For all species tested, the depleted RNA had significantly reduced 18S (Fig. 3A) and 28S (Fig. 3B) rRNA levels as compared to the two other groups (p<0.0001 for all comparisons, Two-Way ANOVA with Tukey’s correction). We also subjected both the input RNA and the depleted RNA to electrophoretic analysis using a Bioanalyzer 2100. For all three species tested, the peak for rRNA (both 18S and 28S typically appear at ~2000nt) is inapparent following the depletion protocol (Fig. 3C-E). In contrast, the input RNA has a prominent peak for rRNA. The traces for the depleted and input RNA are overlaid on the same graph to facilitate comparison.

**Figure 3:**
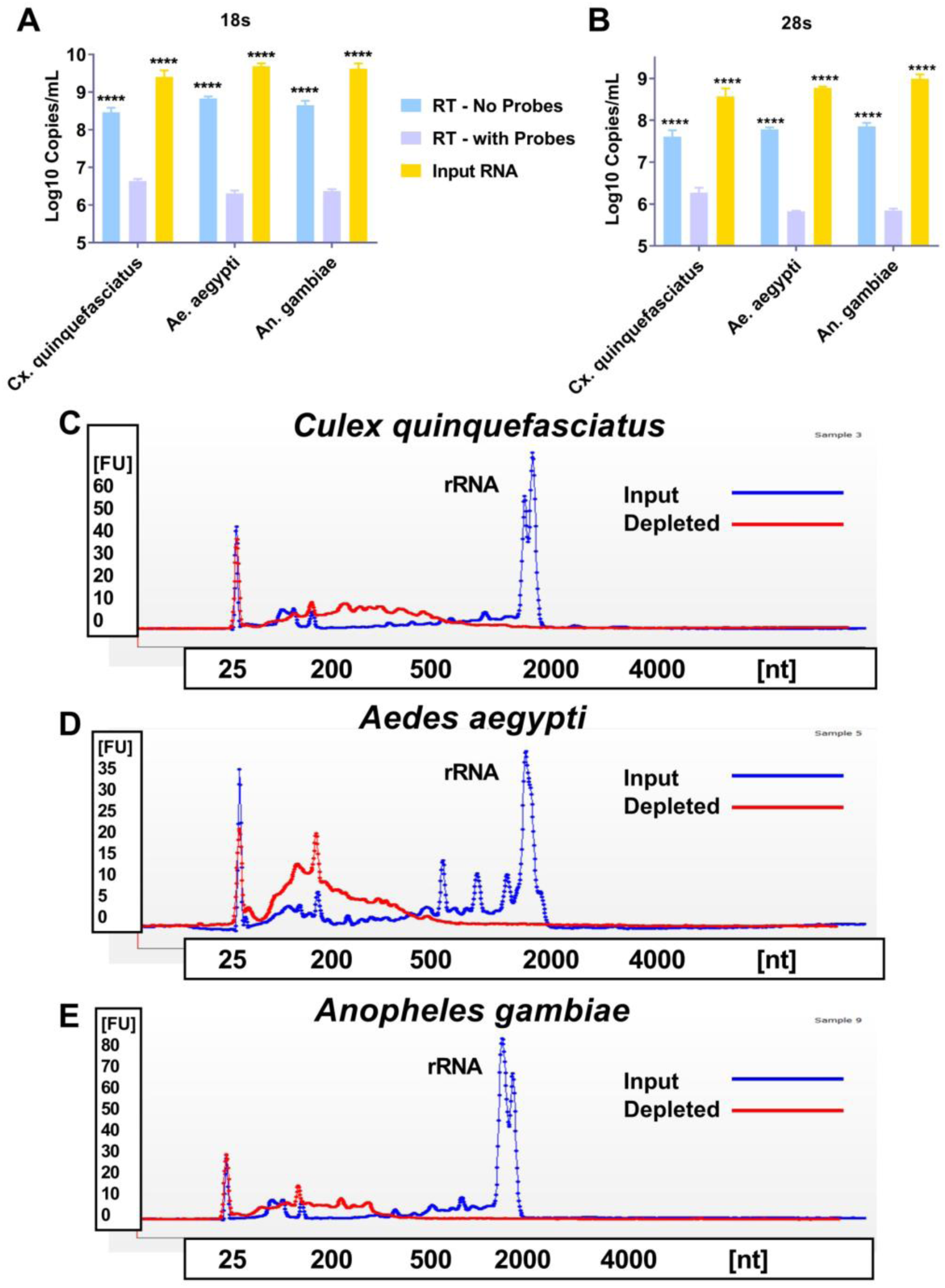
**Reverse Transcriptase mediated ribosomal RNA (rRNA) depletion is effective against mosquitoes from three distinct medically relevant genera.** Total RNA was extracted from three distinct pools of whole mosquitoes from three medically relevant genera; *Culex (Cx.) quinquefaeciatus*, *Aedes (Ae.) aegpyti* and *Anopheles (An.) gambiae.* The RNA was treated with DNase I and then purified; this will now be called Input RNA. An aliquot was then taken and reverse transcribed to cDNA using AMV reverse transcriptase (RT) and DNA probes specific for mosquito ribosomal RNA (RT – with Probes) or in the absence of probes (RT – No Probes). The samples were then treated with RNase H and DNase I to remove the RNA present in an RNA:DNA hybrid and cDNA, respectively. The samples were then purified and subjected to qRT-PCR with primer probe combinations specific for 18S or 28S rRNA (A and B). The Input RNA and RT – with Probes were then assessed using a Bioanalyzer. Panels C-E show a representative trace for each of the three mosquito species tested, *Cx. quinquefaeciatus* (C), *Ae. aegypti* (D), *An. gambiae* (E). The blue trace for each panel shows the Input RNA and the red trace shows the RT – with Probes treated RNA. The peak present at roughly 40 seconds in each trace is the peak for both 18S and 28S rRNA.

### RT mediated rRNA depletion increases sequencing reads to viruses and mRNA

Samples to test depletion efficacy were prepared using a method termed Xenosurveillance, prepared as described previously (Fauver et al., 2018). Briefly, *An. gambiae* mosquitoes were exposed to an infectious bloodmeal containing WNV (which doesn’t replicate in these mosquitoes) and then midguts containing the partially digested blood were collected the next day on FTA cards. The nucleic acids were eluted and extracted as previously described and then the samples were depleted with AMV RT and depletion probes (RT - with Probes). We also tested the Input RNA and samples that underwent the depletion protocol with the omission of probes (RT - No Probes). Following depletion, the RNA underwent Illumina library prep and was sequenced using the MiSeq platform (Illumina). The reads were then trimmed, duplicates removed and mapped to either rRNA (18S (Fig. 4A), 28S (Fig. 4B), host transcriptome (Fig. 4C) or viral (WNV (Fig. 4D), Bolahun virus (BOLV, Fig. 4E) sequences. BOLV is known to persistently infect these mosquitoes (Fauver et al., 2016). A significantly lower proportion of reads mapped to rRNA in the depleted RNA (One-Way ANOVA with Tukey’s correction, p<0.0001 for all comparisons to depleted). In contrast, a significantly increased proportion of the reads aligned to sequences of interest, notably the host transcriptome and the two viruses, WNV and BOLV (One-Way ANOVA with Tukey’s correction, all p<0.01 or lower for all comparisons to depleted). Coverage plots from input, depleted and non-depleted RNA samples are presented in Supplemental Figure 2 for both BOLV and WNV. Finally, we assessed the ability to analyze intrahost viral variation in BOLV by calling variants with LoFreq. A significantly greater number of minority variants could be called in the depleted RNA when compared to the other two groups (One-Way ANOVA with Tukey’s correction, p<0.05 for all comparisons to depleted).

**Figure 4:**
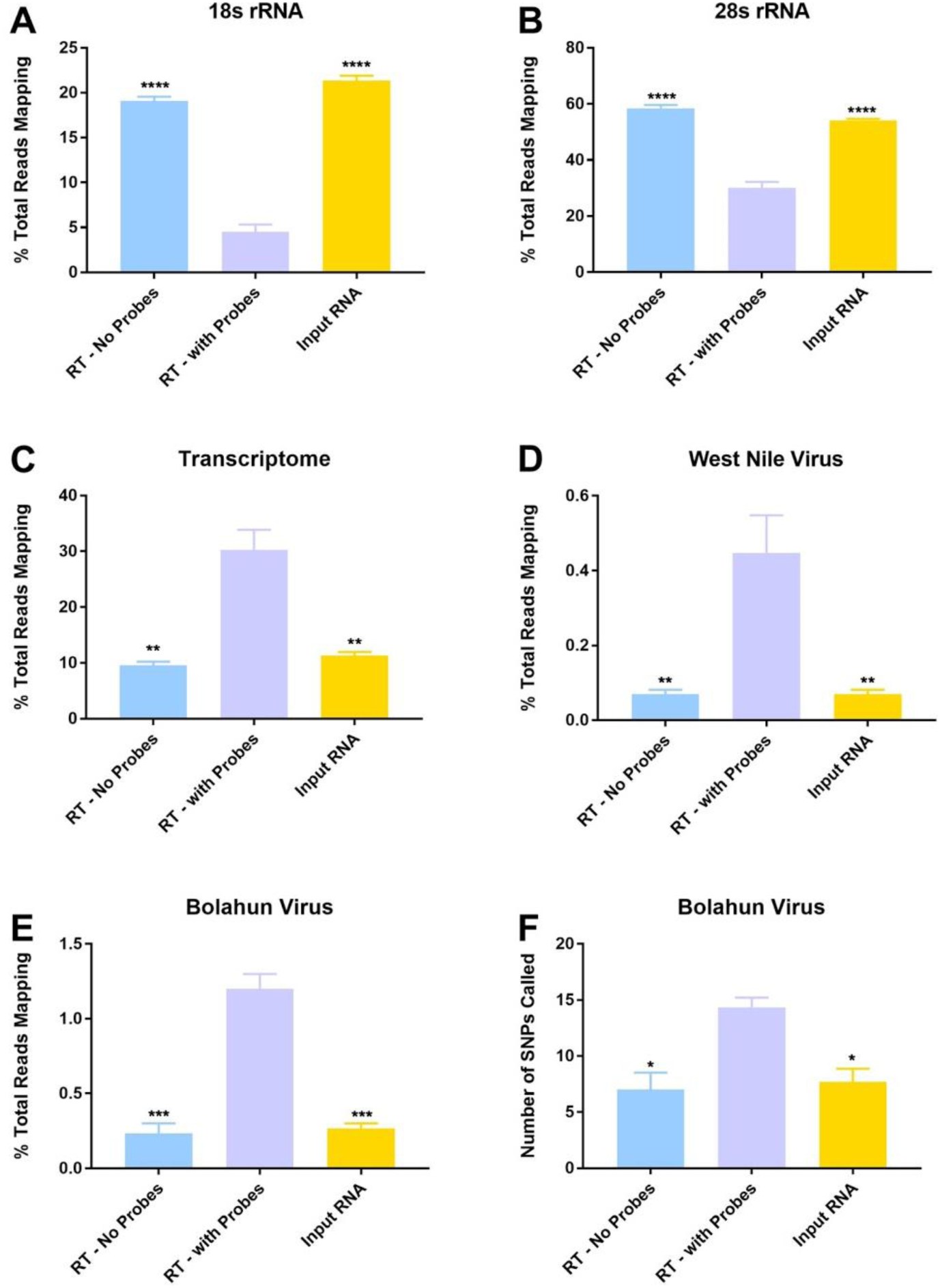
**Reverse Transcriptase mediated ribosomal RNA (rRNA) depletion increases target-specific coverage while reducing the number of rRNA reads in next-generation sequencing.** *Anopheles gambiae* mosquitoes were exposed to an infectious bloodmeal containing 10^7^ PFU of West Nile virus strain NY99. The following day, midguts were dissected and the residual bloodmeal was spread onto a CloneSaver FTA card (GE Healthcare, USA) and then soaked in RNAlater solution to stabilize the nucleic acid and facilitate dispersion. Total nucleic acid was then extracted and DNAse treated. This is considered the Input RNA. DNAse-free RNA was then reverse transcribed using either ribosomal RNA specific probes (RT – with Probes) or without probes (RT – no Probes). The samples were then treated with RNAse H and DNAse I and purified. The samples were then subjected to library preparation and sequenced on an Illumina MiSeq. Reads were then demultiplexed and subsequently trimmed using BBDuk. Duplicate reads were removed using Clumpify and then unique reads were mapped using Bowtie2 to the appropriate reference sequence, 18S rRNA (A), 28S rRNA (B), *An. gambiae* transcriptome (C), West Nile virus (D) and Bolahun virus (E). Percentage of reads mapping was calculated using MultiQC. Variants detected in Bolahun virus were called using LoFreq (F).

### Mosquito collections and sequencing summary

A total of 978 adult field-collected mosquitoes from 10 species were pooled for analysis by NGS (Supplemental Table 2). The most abundant species collected (242) was *Coquillettidia venezuelensis,* followed by *Ae. albopictus* (238), *Psorophora albipes* (110), *Ps. varipes* (101), *Ae. angustivittatus* (91), *Cx. nigripalpus* (87), *Ae. aegypti* (72), *Ae. taeniorhynchus* (33), *Ae. serratus* (2), and *Ps. ferox* (2). All species collected in this study have previously been reported from Chiapas state (Bond et al., 2014; Heinemann and Belkin, 1977). A single MiSeq run following quality filtering and removal of duplicate reads yielded 25.9 million total reads, resulting in 3.8Gb of paired-end data. The total percentage of reads mapping to rRNA in the field samples was in line with what we observed after depletion in our colony mosquitoes (Supplemental Fig. 3). In contrast, the percentage of reads mapping to viruses was relatively high, particularly for *Ae. aegypti*. The percentage of reads mapping to viruses varied widely between the different mosquito species tested.

### Virus sequences identified in field collected mosquitoes following rRNA depletion

Each mosquito species sequenced, save a single pool of 2 *Ps. ferox* mosquitoes, produced contigs aligning to known viral sequences (Fig. 5, Supplemental Table 3). Based off of amino acid similarity and phylogenetic placement, 8 major clades as well as multiple families of RNA viruses were represented across all samples. Amino acid similarities spanned anywhere from 28% (Reovirus contig from *Ae. angustivittatus*) to 99% (Phasi Charoen-like phasivirus RDRP from multiple *Aedes* species). Multiple previously described viruses were identified, based on >95% pairwise nucleotide identity, including Phasi Charoen-like phasivirus (PCLV) in *Ae. aegypti, Ae. angustivittatus,* and *Ps. varipes*. A complete genome of PCLV was assembled from pools of both male and female *Ae. aegypti* mosquitoes (Supplemental Fig. 4). This PCLV genome aligned to Phasi Charoen-like phasivirus strain 2b (Accession: MH237598) with ~98% pairwise nucleotide identity. PCLV sequences from *Ae. angustivittatus* and *Ps. varipes* aligned only to a portion of the RDRP. Partial sequences aligning to both the RDRP and capsid proteins of Humaita-Tubiacanga (HTV) virus were identified from female *Ae. aegypti* and male *Ae. albopictus* mosquitoes. Sequences aligned to HTV with 98.5 and 97.5% pairwise nucleotide identity, respectively.

**Figure 5:**
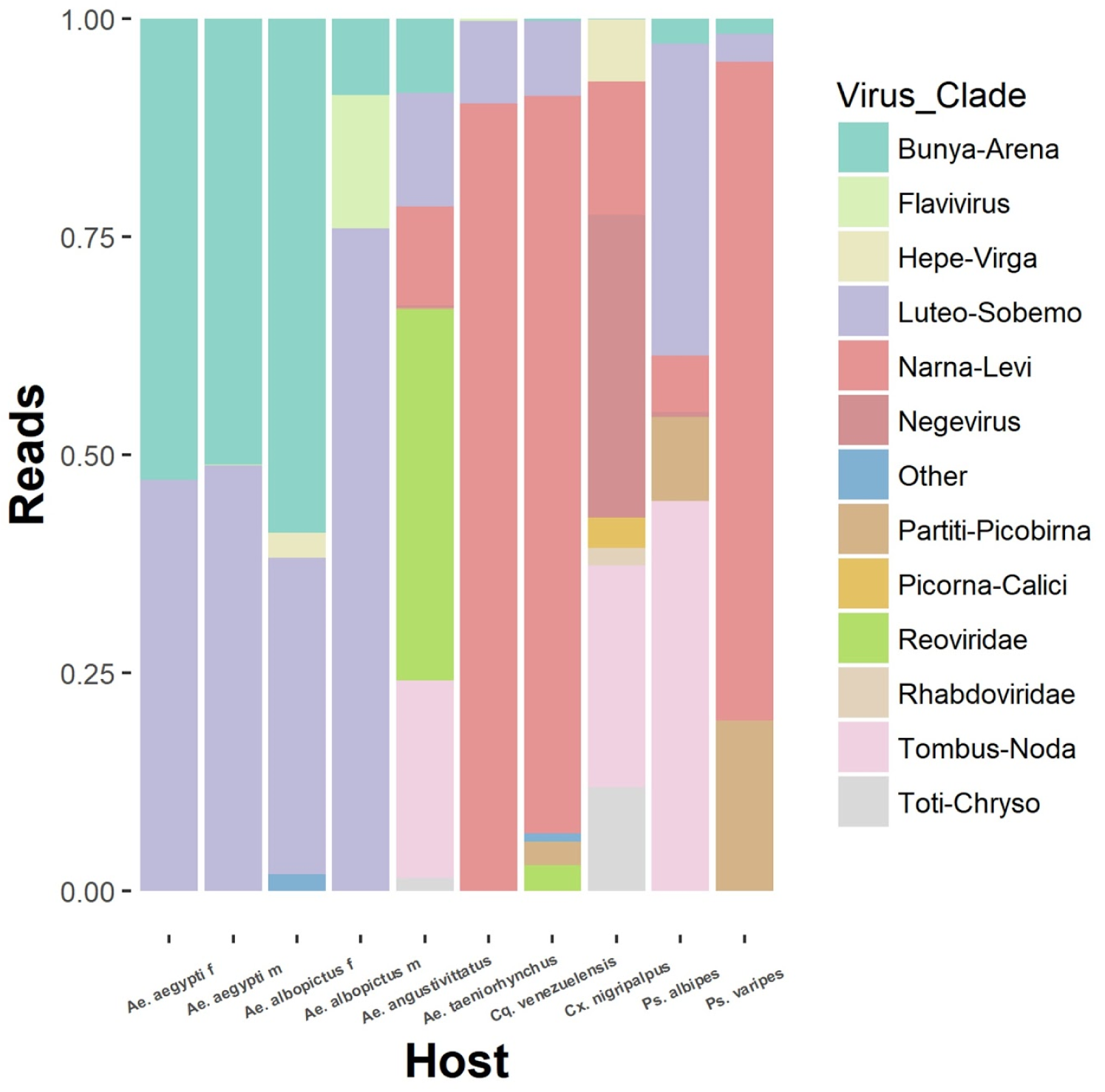
**Viral sequences belonging to diverse clades of RNA viruses identified in field-collected mosquitoes following rRNA depletion.** Individual reads from each mosquito species sequenced were mapped back to all virus contigs identified in this study. Virus clade is inferred by amino acid similarity to other closely related sequences.

Short flavivirus sequences (100-250) were found in 7 of 8 mosquito species sequenced aligning to the same portion of the West Nile virus (WNV) genome. Based on the sequence similarity between species, its presence in nearly all groups, and our frequent use of WNV in our laboratory, it is likely these sequences are the result of laboratory contamination during library preparation opposed to an authentic infection in our mosquito samples.

While numerous contigs were generated that distantly resembled known viral sequences, indicating the presence of divergent viruses in these species, we chose to further analyze only contigs that produced coding complete viral genomes (Ladner et al., 2014). Our computational approached generated 6 novel viral genomes, including a novel strain of a previously described negevirus (Fig. 6A), 5 Levi-Narnaviruses (Fig. 7A-E), and 1 Luteo-Sobemo virus (Fig. 8A).

**Figure 6:**
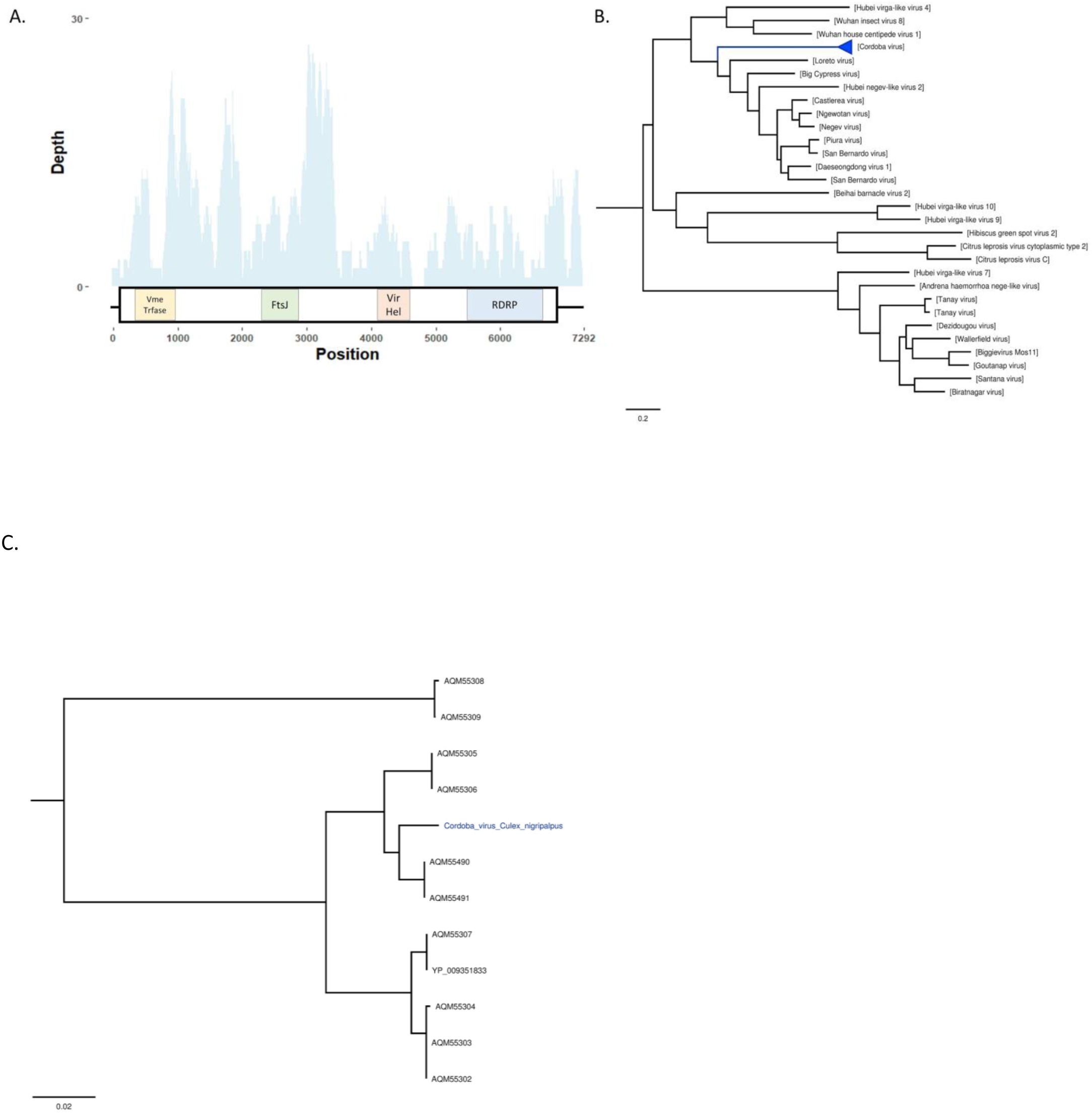
**Description of a novel variant of the negevirus Cordoba virus from *Culex nigripalpus*** A- Virus cartoon depicting the genomic structure and depth of coverage Cordoba virus *Cx. nigripalpus* variant. The large boxes represent predicted ORFs and the small boxes represent areas of protein homology to viral methyltransferase (pfam01660), FtsJ-like methyltransferase (pfam01728), viral RNA helicase (pfam01443), and viral RNA-dependent RNA polymerases (cd1699). B. Phylogenetic placement of multiple strains of Cordoba virus highlighted in blue. Phylogenies were created using 1,234 A.A. residues across the complete ORF. Tree is midpoint rooted. Phylogenetic trees were generated in FigTree. C. Expansion of phylogenetic tree containing the sequenced strains of Cordoba virus. The strain sequenced in this study is highlighted in blue. Phylogenetic trees were generated in FigTree.

**Figure 7:**
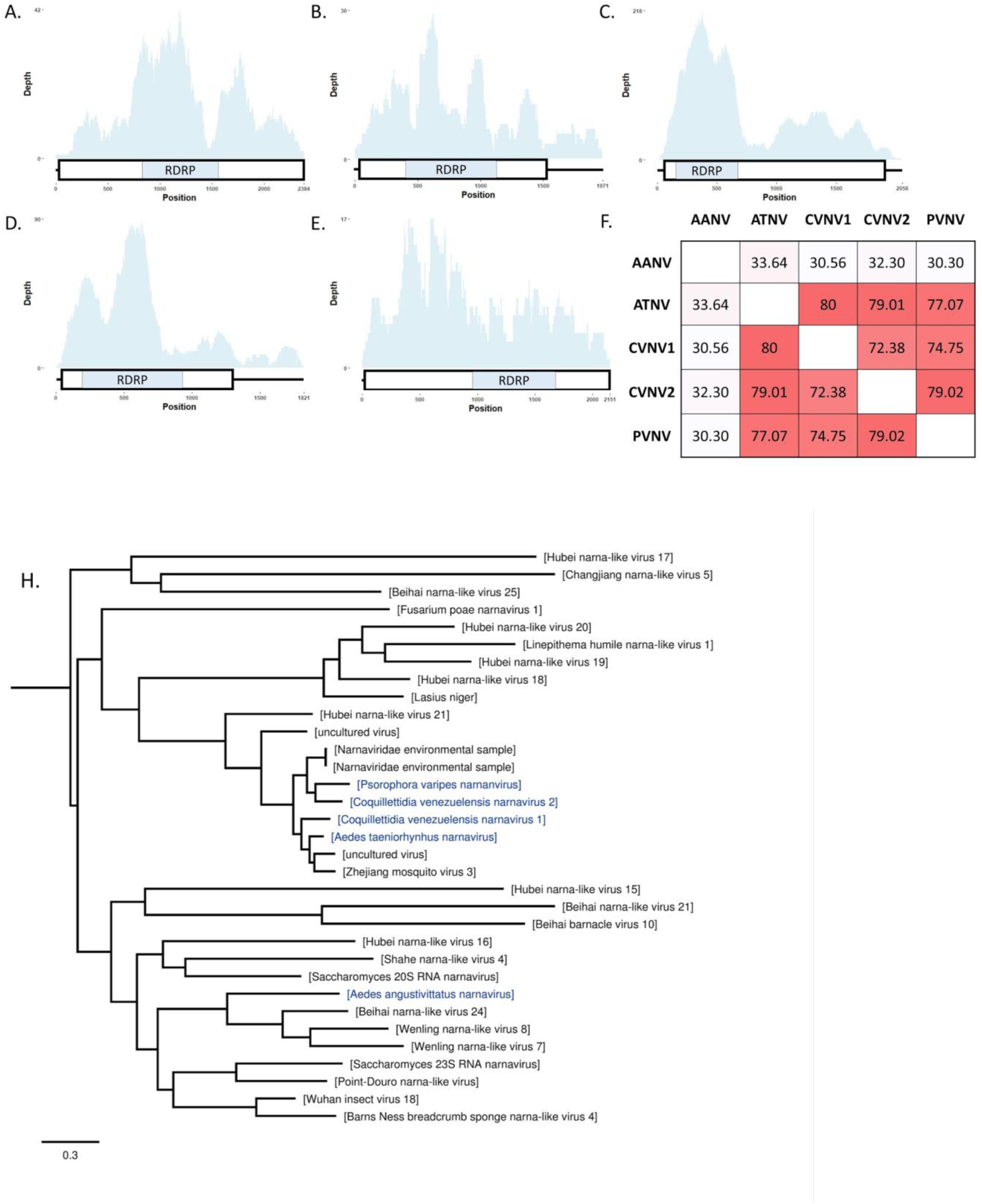
**Multiple, unique narnaviruses described from multiple mosquito species.** A-E cartoons depicting the simple genomic structure and depth of read coverage to newly described narnaviruses. The large boxes represent predicted ORFs and the small boxes represent protein homology to viral RNA-dependent RNA polymerases (cd1699). A- CVNV1, B- CVNV2, C- PVNV, D- ATNV, E- AANV. F- Pairwise identify of 295 amino acid residues across the predicted RDRP between the newly described narnaviruses. H- Phylogenetic placement of novel narnaviruses highlighted in blue. Tree based on alignments of RDRP from multiple narnavirus and is midpoint rooted. Phylogenetic trees were generated in FigTree.

**Figure 8:**
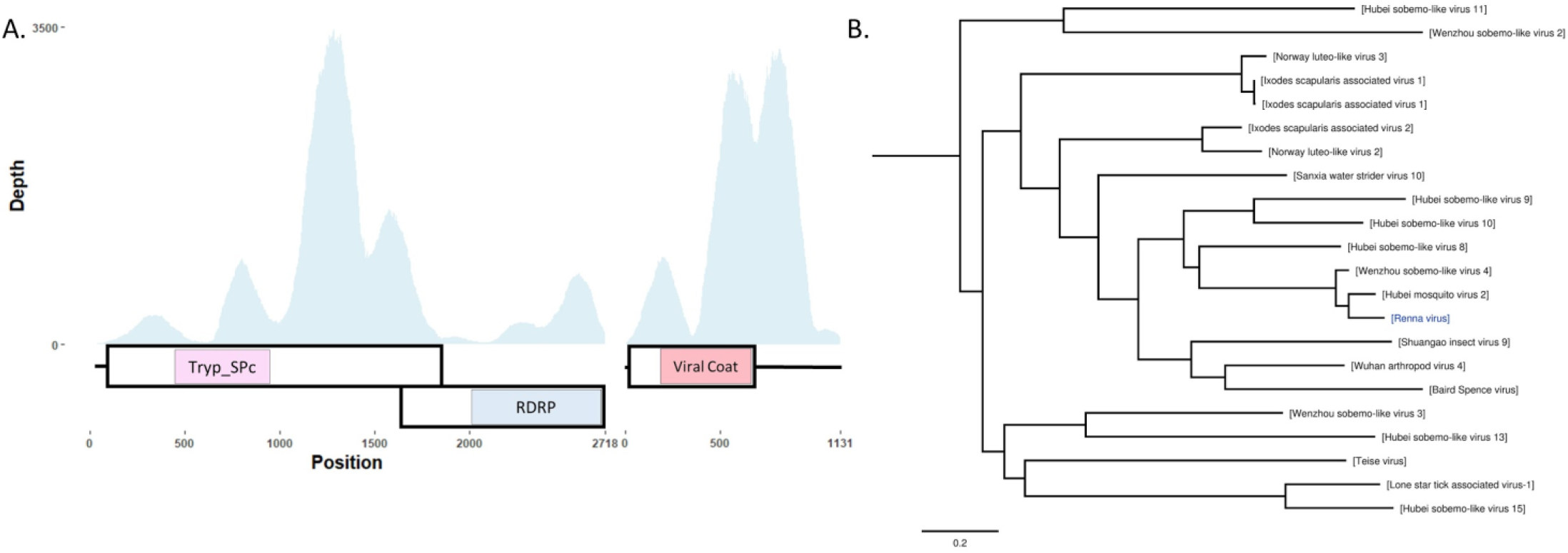
**Description of a novel Luteo-Sobemo like virus from *Aedes aegypti* mosquitoes.** A- Cartoon depicting the predicted bipartite genomic structure of RENV. Large boxes represent ORFs, small boxes represent areas of protein homology to Trypsin-like serine protease (cd00190), viral RNA-dependent RNA polymerase (cd1699), and capsid protein (cd00205). B- Phlyogenetic placement of RENV. Phylogeny was created using a 289 amino acid portion of the RDRP. Trees are midpoint rooted. Phylogenetic trees were generated in FigTree.

A total of 4 contigs identified in *Cx. nigripalpus* mosquitoes aligned to the CoB_37B strain of Cordoba virus with estimated gaps of 188, 72, and 55 nucleotides. The assembly of these contigs produced a final sequence approximately 7,300 nucleotides long that contained a single ORF predicted to code for 4 proteins (Fig. 6A). These proteins include a viral methyltransferase (pfam01660), FtsJ-like methyltransferase (pfam01728), Viral RNA helicase (pfam01443), and RDRP (cd01699) (Fig. 6A). Both the type of proteins encoded and synteny of the genome are in agreement with representative +ssRNA viruses from the Nelorpivirus group of Negeviruses (Nunes et al., 2017). Phylogenetic placement and high pairwise nucleotide identity (78.8-93.6%, depending on strain) indicated this genome to be a novel strain of Cordoba virus, a negevirus described from a variety of mosquito species, including *Cx. nigripalpus*, from Nepal, the U.S., and Colombia (Nunes et al., 2017) (Fig. 6B,C).

Multiple sequences related to viruses in the +ssRNA Narna-Levi clade were identified from *Ae. angustivittatus*, *Ae. taeniorhynchus*, *Cq. venezuelensis*, and *Ps. varipes*. Two distinct contigs were generated from *Cq. venezuelensis* mosquitoes. These sequences were found to be approximately 2kb in length and contain a single ORF that encodes for RDRP (cd01699) (Fig. 7 A-E). Pairwise amino acid identity was approximately 72-80% between 4 of the virus sequences, while a sequence from pools of *Ae. angustivittatus* mosquitoes varied substantially (30-33%) compared to other sequences described in this study (Fig. 7F). The 4 more similar genomes grouped with other narnavirus-like sequences described from mosquitoes, where the sequence from *Ae. angustivittatus* mosquitoes grouped with narnavirus-like sequences from crustaceans (Fig. 7H). These virus genomes have provisionally been designated Aedes angustivittatus narnavirus (AANV), Aedes taeniorhynchus narnavirus (ATNV), Coquillettidia venezuelensis naranvirus 1 & 2 (CVNV1, CVNV2), and Psorophora varipes narnavirus (PVNV).

Two sequences related to +ssRNA Luteo-Sobemo like viruses, 2,718 and 1,131 nucleotides in length, were identified in pools of both male and female *Ae. aegypti* mosquitoes. The longer sequence is predicted to encode for two proteins, a Trypsin-like serine protease (cd00190) and RDRP (cd01699), respectively, in two separate ORFs (Fig. 8A). These ORFs overlap and appear to be on the same segment indicating the reading frame difference is likely the result of frameshift mutation, which is common in Luteo-Sobemo viruses (Barry and Miller 2002). The identified “slippery sequence”, a conserved heptanucleotide sequence that causes the ribosome to shift reading frames, in Sobemoviruses is “UUUAAAC”(Mäkinen et al. 1995). This specific sequence was not identified, however, as these viruses are divergent and not well characterized, it is possible a non-canonical heptanucleotide sequence could exist. A sequence 24 base pairs upstream of the second ORF reads “GGGCCCG”, which deviates slightly from the typical slippery sequence construct of “XXXYYYZ” (P. 2012). It remains to be determined whether this sequence is responsible for ribosomal frameshifting in this virus. The smaller sequence contains a single ORF encoding the predicted viral coat protein (pfam00729). The bipartite genomic structure is seen in a similar virus, Hubei mosquito virus 2 (Shi et al., 2016). This sequence, provisionally named Renna virus (RENV), groups phylogenetically with viruses identified from a variety of ticks and insects, including mosquitoes (Fig. 8B). Both segments had a high average depth of coverage, 650 and 1351, respectively in *Ae. aegypti* females. RENV from male and female *Ae. aegypti* mosquito pools shared a >99% pairwise nucleotide identity.

## Discussion

Studies involving sequencing viral RNA; such as viral metagenomics, intrahost viral dynamics, transcriptomics and virus discovery require target reads to be at sufficient levels to perform meaningful analysis. These analyses are often hampered by the high percentage of ribosomal RNA (rRNA) present in total RNA, which can reach greater than 80-90% of the total sample (Eun, 1996). Since these reads are rarely used, this represents significant waste of both financial and computational resources, and limits the amount of multiplexing that can be performed. While procedures such as selection of polyadenylated transcripts can be used to enrich RNA preparations for mRNA, this is not relevant to RNA viruses that lack polyadenylation. Furthermore, other methods like amplicon sequencing or probe capture are sequence specific, and thus unknown pathogens cannot be sampled. Therefore, selective depletion of highly abundant rRNA is beneficial. Several methods and commercial kits are available to do this but most are designed to work specifically for human or mouse samples. Here, we describe a novel method that utilizes specific reverse transcription of rRNA using small DNA probes for depletion along with RNase H. This allowed us to design depletion probes that could simultaneously deplete rRNA from mosquitoes of highly diverse genetic backgrounds. Using this method, we show that specific depletion of rRNA results in increased reads to meaningful RNA, such as viruses and host mRNA. In addition, we detected more intrahost variants using this depletion method. Although we subjected all field-collected mosquito pools to rRNA depletion, thus no non-depleted libraries were generated for comparison, we were able to detect novel virus genomes from a single, highly multiplexed (64 libraries), MiSeq run of nearly 1,000 diverse field-collected mosquito samples that underwent rRNA depletion. Taken together, these findings suggest that RT-mediated rRNA depletion can facilitate sequencing sequencing of mosquito samples both from the lab and field.

To our knowledge, only two other studies have aimed to assess rRNA depletion strategies from insect species. The first used a commercial kit designed for mammalian rRNA, Epicentre’s Ribo-Zero rRNA, to deplete rRNA from *Drosophila* flies. While the approach seemed to effectively remove rRNA and enrich mRNA transcripts, it suffers from being high-cost (Kumar et al., 2012). Another study showed by bioanalyzer and NGS effective removal of rRNA from mosquito midguts using RNA probes to the rRNA (Kukutla et al., 2013). However, this technique required large amounts of input RNA (50 pooled midguts), uses unstable RNA probes and expensive streptavidin beads. Furthermore, it’s unclear if this technique works for other species or just *An. gambiae.* Accordingly, we devised a novel method for depleting rRNA using RNase H depletion that was based on the method described by Morlan et al. (Morlan et al., 2012) with the exception that it uses shorter probes and incorporates a reverse transcription (RT) step. The shorter probes allow highly conserved regions to be targeted, thus making it possible to simultaneously deplete rRNA from high divergent species or even genera. The RT step extends the bound DNA probes to produce cDNA complementary to the rRNA that is destroyed following both a RNase H and DNase I digestion.

First, we assessed the efficacy of several RTs to convert rRNA to cDNA and subsequently be degraded by RNase H. We found AMV to be the optimal enzyme, depleting a significant amount of rRNA with no off-target effects. While M-MLV RT depleted as much or more rRNA as AMV, it also depleted WNV RNA, suggesting it was converting non-target RNA species to cDNA as well (Fig. 2C). It’s unclear why M-MLV RT would have off-target effects and not AMV, especially as there has been evidence of the opposite occuring in a previous publication (Agranovsky, 1992). This and other publications have shown primer-independent cDNA synthesis for both AMV and M-MLV RTs, which could explain the high level of non-specific depletion in M-MLV but not AMV observed here (Freeh and Peterhans, 1994). Agranovsky et al. presented evidence that a tRNA contaminant in the AMV RT preparation tested at that time was responsible for this primer-independent cDNA synthesis. We cannot rule out the possibility that the M-MLV obtained from NEB contained some contaminant that could effectively primer non-target RNA species such as WNV. There may also be small RNAs present in our samples that could have primed cDNA synthesis particularly well for M-MLV. Both Superscript (SS) III and IV, mutants of M-MLV were effective at depleting 18S rRNA with no off-target effects. While SSIII also depleted 28S rRNA, SSIV did not effectively deplete this RNA species. Finally, Tth DNA polymerase, which shows RT activity in the presence of manganese, did not effectively deplete rRNA, even in the presence of specific DNA probes. This may be related to the fact that Tth and SSIV lack functional RNase H domains (Myers and Gelfand, 1991), suggesting that this intrinsic activity is important for the mechanism of depletion with this technique, even if RNase H is added after the RT step. Next, we assessed whether the RT step was necessary for depletion in the workflow, as Morlan et al. had previously shown efficient depletion in the absence of this step (Morlan et al., 2012). The RT step was critical to the depletion observed and the specific depletion probes were necessary, as samples treated with DNA probes in the absence of RT had only a modest depletion effect. However, in the presence of RT and specific depletion probes, 18S and 28S rRNA were depleted roughly 100- and 1000-fold, respectively.

Depletion was then tested on RNA from three medically important mosquito species representing three distinct genera; *Ae. aegypti, An. gambiae* and *Cx. quinquefasciatus*. These species transmit a significant proportion of vector-borne pathogens; including dengue virus, Zika virus, chikungunya virus, malaria parasites and West Nile virus, among others. We found by qRT-PCR and bioanalyzer, depletion in the presence of rRNA probes was associated with a significant reduction in 18S and 28S rRNA from all three species tested. Despite the almost complete removal of the peak for rRNA in the bioanalyzer traces, we were still able to detect rRNA sequences by both qRT-PCR and NGS. This might be a result of incomplete digestion of the RNA by RNase H due to incomplete activity or RNA that hadn’t been reverse transcribed. It’s possible that the secondary structure of rRNA prevents the complete synthesis of cDNA from RNA and that this is not degraded by the RNase H. Different methods to increase the efficiency of cDNA synthesis or adding additional DNA probes may be beneficial in future iterations of this protocol. This result suggested that this protocol could be used for a wide array of mosquito species, as *Aedes* and *Culex* are significantly divergent from *Anopheles* mosquitoes, having separated likely over 200 million years ago (Reidenbach et al., 2009). In fact, we have seen rRNA depletion by NGS in virus stocks prepared in mammalian cells as well, suggesting a broad range of cross-reactivity to rRNA from different species.

We then depleted rRNA from midguts isolated from *An. gambiae* mosquitoes that were fed a bloodmeal containing WNV. This RNA was then subjected to Illumina deep-sequencing and the resulting reads were aligned to several sequences. We observed significant depletion of rRNA while increasing the percentage of reads to mRNA, WNV and the insect-specific virus Bolahun virus (Fauver et al., 2016). We were also able to identify significantly more minority variants present in Bolahun virus, suggesting intrahost virus population analyses are facilitated following depletion. It has been shown that high levels of sequencing coverage are necessary to perform intrahost virus analysis, which is can be difficult to achieve without depletion or enrichment (McCrone and Lauring, 2016).

As second and third generation sequencing based approaches for the detection and analysis of vector-borne pathogens from field-collected mosquitoes are becoming commonplace, techniques that increase reads to target sequences in complex samples will be sorely needed. Accordingly, we employed our rRNA depletion method to a diverse group of field-collected mosquitoes and subjected them to NGS with the goal of identifying both human-infecting and insect-specific viruses. While we did not identify arbovirus sequences from these pools of mosquitoes, we were able to identify partial and coding complete genomic sequences of a variety of presumed insect specific viruses. As all pools were subjected to rRNA depletion, we do not have non-depleted libraries to compare the efficacy of rRNA depletion to. However, the total number of reads aligning to rRNA from these samples was congruent with what we observed in our laboratory studies. In fact, in libraries constructed from *Ae. aegypti* females, more reads competitively aligned to viruses than to 28s or 18s rRNA sequences, although the number of reads aligning to both viruses and rRNA sequences varied widely between divergent genera. Using our bioinformatic approach, 7 novel coding complete viral genomes were identified, in addition to the previously described insect specific viruses PCLV and HTV. Complete PCLV genomes were assembled from pools of both male and female *Ae. aegypti* mosquitoes at a relatively high depth of coverage and pairwise nucleotide identity. PCLV has been identified mosquito cell culture and in numerous populations of *Ae. aegypti* mosquitoes from across the globe (Chandler et al., 2014; Di Giallonardo et al., 2018; Yamao et al., 2009; Zhang et al., 2018). In addition to PCLV, we identified large contigs with >99% nucleotide identity to HTV in both female *Ae. aegypti* and male *Ae. albopictus* mosquitoes (Aguiar et al., 2015; Zakrzewski et al., 2018). A total of 5 coding complete narnavirus genome sequences were identified from 4 species of mosquitoes collected in this study. Of the 5 virus genomes described here, 4 group closely together and with other narnaviruses described from mosquitoes. While multiple narnaviruses have been identified by metagenomic sequencing of whole mosquito samples, it remains to be determined if these represent infections of fungi in the normal microbiota, or bona fide infections of mosquitoes (Chandler et al., 2015; Cook et al., 2013; Shi et al., 2016). A novel strain of Cordoba virus, a negevirus described previously from mosquitoes, was identified in *Cx. nigripalpus* mosquitoes (Nunes et al., 2017). We were also able to assemble the coding complete genome of RENV, a virus that groups with Luteo-Sobemo viruses identified in mosquitoes (Shi et al., 2016). Based on the phylogenetic placement of these sequences, all the viruses described in this study are presumed to be insect specific, however this is yet to be validated. As well, the effect these viruses may have on mosquito biology or vector competence remains to be determined. It is highly probable that these viruses would have been detected if we did not perform rRNA depletion, but based on our NGS data from laboratory experiments, our depletion method likely aided in discovery and characterization by allowing more unique, non-rRNA sequences to be identified. Although the amount of viral RNA from any given mosquito depends upon individual infection status and the amount of viral replication occurring, we were able to identify and assemble multiple viral genomes from a highly multiplexed sequencing run on a comparatively low-output sequencing platform. Increasing reads to target sequences of interest (e.g. viruses) by depleting uninformative rRNA sequences from complex, field-collected mosquito samples has the potential to improve the efficacy and feasibility of using metagenomic sequencing for mosquito-borne disease surveillance.

## Acknowledgements

We would like to acknowledge Nunya Chotiwan, Rushika Pereira, Karla Saavedra, Ildefonso Fernández-Salas and all of the employees at the Centro Regional de Investigación en Salud Pública for assistance in collecting mosquitoes in Tapachula, Chiapas, Mexico. We also acknowledge the funding sources for providing the resources to perform this work; NIAIDAI067380.

## Supplemental Material

**Supplemental File 1:** Detailed protocol for rRNA depletion.

**Supplemental Figure 1: Position of DNA probes across the mosquito 45S Ribosomal RNA (rRNA) sequence.** The DNA probes were aligned to the 45S rRNA sequence of *Aedes albopictus*. The probes are presented in green and the rRNA sequence is labelled and in red.

**Supplemental Figure 2: Depth of coverage by nucleotide position for Bolahun virus and West Nile virus.** Depth was calculated at each nucleotide position using Samtools depth -a. Three samples were sequenced per group. Grey shading represents the minimum and maximum coverage at each position.

**Supplemental Figure 3: Proportion of total reads mapping to virus, 18s, and 28s sequences from field-collected mosquitoes that have undergone rRNA depletion.**

**Supplemental Figure 4: Depth of coverage of complete Phasi-Chareon like phasivirus from female *Ae. aegypti* mosquitoes.**

**Supplemental Table 1:** List of oligonucleotide probe sequences aligning to mosquito 45S rRNA used for depletion.

**Supplemental Table 2:** Metadata for each library constructed from field-collected mosquitoes.

**Supplemental Table 3:** Description of viral contigs >500 nucleotides in length identified from field-collected mosquitoes.

## References

Adiconis, X., Borges-Rivera, D., Satija, R., DeLuca, D.S., Busby, M.A., Berlin, A.M., Sivachenko, A., Thompson, D.A., Wysoker, A., Fennell, T., Gnirke, A., Pochet, N., Regev, A., Levin, J.Z., 2013. Comparative analysis of RNA sequencing methods for degraded or low-input samples. Nat. Methods 10, 623–629.

Agranovsky, A.A., 1992. Exogenous primer-independent cDNA synthesis with commercial reverse transcriptase preparations on plant virus RNA templates. Anal. Biochem. 203, 163–165.

Aguiar, E.R.G.R., Olmo, R.P., Paro, S., Ferreira, F.V., de Faria, I.J. da S., Todjro, Y.M.H., Lobo, F.P., Kroon, E.G., Meignin, C., Gatherer, D., Imler, J.-L., Marques, J.T., 2015. Sequence-independent characterization of viruses based on the pattern of viral small RNAs produced by the host. Nucleic Acids Res. 43, 6191–6206.

Altschul, S.F., Gish, W., Miller, W., Myers, E.W., Lipman, D.J., 1990. Basic local alignment search tool. J. Mol. Biol. 215, 403–410.

Aubry, M., Teissier, A., Huart, M., Merceron, S., Vanhomwegen, J., Roche, C., Vial, A.-L., Teururai, S., Sicard, S., Paulous, S., Desprès, P., Manuguerra, J.-C., Mallet, H.-P., Musso, D., Deparis, X., Cao-Lormeau, V.-M., 2017. Zika Virus Seroprevalence, French Polynesia, 2014-2015. Emerg. Infect. Dis. 23, 669–672.

Bond, J.G., Casas-Martínez, M., Quiroz-Martínez, H., Novelo-Gutiérrez, R., Marina, C.F., Ulloa, A., Orozco-Bonilla, A., Muñoz, M., Williams, T., 2014. Diversity of mosquitoes and the aquatic insects associated with their oviposition sites along the Pacific coast of Mexico. Parasit. Vectors 7, 41.

Capella-Gutiérrez, S., Silla-Martínez, J.M., Gabaldón, T., 2009. trimAl: a tool for automated alignment trimming in large-scale phylogenetic analyses. Bioinformatics 25, 1972–1973.

Chandler, J.A., Liu, R.M., Bennett, S.N., 2015. RNA shotgun metagenomic sequencing of northern California (USA) mosquitoes uncovers viruses, bacteria, and fungi. Front. Microbiol. 6, 185.

Chandler, J.A., Thongsripong, P., Green, A., Kittayapong, P., Wilcox, B.A., Schroth, G.P., Kapan, D.D., Bennett, S.N., 2014. Metagenomic shotgun sequencing of a Bunyavirus in wild-caught Aedes aegypti from Thailand informs the evolutionary and genomic history of the Phleboviruses. Virology 464-465, 312–319.

Cook, S., Chung, B.Y.-W., Bass, D., Moureau, G., Tang, S., McAlister, E., Culverwell, C.L., Glücksman, E., Wang, H., Brown, T.D.K., Gould, E.A., Harbach, R.E., de Lamballerie, X., Firth, A.E., 2013. Novel virus discovery and genome reconstruction from field RNA samples reveals highly divergent viruses in dipteran hosts. PLoS One 8, e80720.

Cross, S.T., Kapuscinski, M.L., Perino, J., Maertens, B.L., Weger-Lucarelli, J., Ebel, G.D., Stenglein, M.D., 2018. Co-Infection Patterns in Individual Ixodes scapularis Ticks Reveal Associations between Viral, Eukaryotic and Bacterial Microorganisms. Viruses 10. https://doi.org/10.3390/v10070388

Darsie, R.F., Ward, R.A., n.d. Identification and geographical distribution of the mosquitoes of North America, north of Mexico. 2005. Gainesville: University Press of Florida Google Scholar.

Di Giallonardo, F., Audsley, M.D., Shi, M., Young, P.R., McGraw, E.A., Holmes, E.C., 2018. Complete genome of Aedes aegypti anphevirus in the Aag2 mosquito cell line. J. Gen. Virol. 99, 832–836.

Eun, H.-M., 1996. Enzymology Primer for Recombinant DNA Technology. Elsevier.

Fauver, J.R., Grubaugh, N.D., Krajacich, B.J., Weger-Lucarelli, J., Lakin, S.M., Fakoli, L.S., III, Bolay, F.K., Diclaro, J.W., II, Dabiré, K.R., Foy, B.D., Others, 2016. West African Anopheles gambiae mosquitoes harbor a taxonomically diverse virome including new insect-specific flaviviruses, mononegaviruses, and totiviruses. Virology 498, 288–299.

Fauver, J.R., Weger-Lucarelli, J., Fakoli, L.S., 3rd, Bolay, K., Bolay, F.K., Diclaro, J.W., 2nd, Brackney, D.E., Foy, B.D., Stenglein, M.D., Ebel, G.D., 2018. Xenosurveillance reflects traditional sampling techniques for the identification of human pathogens: A comparative study in West Africa. PLoS Negl. Trop. Dis. 12, e0006348.

Forni, D., Filippi, G., Cagliani, R., De Gioia, L., Pozzoli, U., Al-Daghri, N., Clerici, M., Sironi, M., 2015. The heptad repeat region is a major selection target in MERS-CoV and related coronaviruses. Sci Rep5: 14480.

Freeh, B., Peterhans, E., 1994. RT-PCR: “background priming” during reverse transcription. Nucleic Acids Res. 22, 4342–4343.

Grubaugh, N.D., Weger-Lucarelli, J., Murrieta, R.A., Fauver, J.R., Garcia-Luna, S.M., Prasad, A.N., Black, W.C., 4th, Ebel, G.D., 2016. Genetic Drift during Systemic Arbovirus Infection of Mosquito Vectors Leads to Decreased Relative Fitness during Host Switching. Cell Host Microbe 19, 481–492.

Guindon, S., Dufayard, J.-F., Lefort, V., Anisimova, M., Hordijk, W., Gascuel, O., 2010. New algorithms and methods to estimate maximum-likelihood phylogenies: assessing the performance of PhyML 3.0. Syst. Biol. 59, 307–321.

Heinemann, S.J., Belkin, J.N., 1977. Collection records of the project “Mosquitoes of Middle America” 9. Mexico (MEX, MF, MT, MX). Mosq. Syst 9, 483–535.

Jones, K.E., Patel, N.G., Levy, M.A., Storeygard, A., Balk, D., Gittleman, J.L., Daszak, P., 2008. Global trends in emerging infectious diseases. Nature 451, 990–993.

Katoh, K., Standley, D.M., 2013. MAFFT multiple sequence alignment software version 7: improvements in performance and usability. Mol. Biol. Evol. 30, 772–780.

Kearse, M., Moir, R., Wilson, A., Stones-Havas, S., Cheung, M., Sturrock, S., Buxton, S., Cooper, A., Markowitz, S., Duran, C., Thierer, T., Ashton, B., Meintjes, P., Drummond, A., 2012. Geneious Basic: an integrated and extendable desktop software platform for the organization and analysis of sequence data. Bioinformatics 28, 1647–1649.

Kukutla, P., Steritz, M., Xu, J., 2013. Depletion of ribosomal RNA for mosquito gut metagenomic RNA-seq. J. Vis. Exp. https://doi.org/10.3791/50093

Kumar, N., Creasy, T., Sun, Y., Flowers, M., Tallon, L.J., Dunning Hotopp, J.C., 2012. Efficient subtraction of insect rRNA prior to transcriptome analysis of Wolbachia-Drosophila lateral gene transfer. BMC Res. Notes 5, 230.

Ladner, J.T., Beitzel, B., Chain, P.S.G., Davenport, M.G., Donaldson, E.F., Frieman, M., Kugelman, J.R., Kuhn, J.H., O’Rear, J., Sabeti, P.C., Wentworth, D.E., Wiley, M.R., Yu, G.-Y., Threat Characterization Consortium, Sozhamannan, S., Bradburne, C., Palacios, G., 2014. Standards for sequencing viral genomes in the era of high-throughput sequencing. MBio 5, e01360–14.

Lanciotti, R.S., Kerst, A.J., Nasci, R.S., Godsey, M.S., Mitchell, C.J., Savage, H.M., Komar, N., Panella, N.A., Allen, B.C., Volpe, K.E., Davis, B.S., Roehrig, J.T., 2000. Rapid detection of west nile virus from human clinical specimens, field-collected mosquitoes, and avian samples by a TaqMan reverse transcriptase-PCR assay. J. Clin. Microbiol. 38, 4066–4071.

Langmead, B., Salzberg, S.L., 2012. Fast gapped-read alignment with Bowtie 2. Nat. Methods 9, 357–359.

Li, H., Handsaker, B., Wysoker, A., Fennell, T., Ruan, J., Homer, N., Marth, G., Abecasis, G., Durbin, R., 1000 Genome Project Data Processing Subgroup, 2009. The Sequence Alignment/Map format and SAMtools. Bioinformatics 25, 2078–2079.

Li, W., Godzik, A., 2006. Cd-hit: a fast program for clustering and comparing large sets of protein or nucleotide sequences. Bioinformatics 22, 1658–1659.

Matranga, C.B., Gladden-Young, A., Qu, J., Winnicki, S., Nosamiefan, D., Levin, J.Z., Sabeti, P.C., 2016. Unbiased Deep Sequencing of RNA Viruses from Clinical Samples. J. Vis. Exp. https://doi.org/10.3791/54117

McCrone, J.T., Lauring, A.S., 2016. Measurements of Intrahost Viral Diversity Are Extremely Sensitive to Systematic Errors in Variant Calling. J. Virol. 90, 6884–6895.

Metsky, H.C., Matranga, C.B., Wohl, S., Schaffner, S.F., Freije, C.A., Winnicki, S.M., West, K., Qu, J., Baniecki, M.L., Gladden-Young, A., Lin, A.E., Tomkins-Tinch, C.H., Ye, S.H., Park, D.J., Luo, C.Y., Barnes, K.G., Shah, R.R., Chak, B., Barbosa-Lima, G., Delatorre, E., Vieira, Y.R., Paul, L.M., Tan, A.L., Barcellona, C.M., Porcelli, M.C., Vasquez, C., Cannons, A.C., Cone, M.R., Hogan, K.N., Kopp, E.W., Anzinger, J.J., Garcia, K.F., Parham, L.A., Ramírez, R.M.G., Montoya, M.C.M., Rojas, D.P., Brown, C.M., Hennigan, S., Sabina, B., Scotland, S., Gangavarapu, K., Grubaugh, N.D., Oliveira, G., Robles-Sikisaka, R., Rambaut, A., Gehrke, L., Smole, S., Halloran, M.E., Villar, L., Mattar, S., Lorenzana, I., Cerbino-Neto, J., Valim, C., Degrave, W., Bozza, P.T., Gnirke, A., Andersen, K.G., Isern, S., Michael, S.F., Bozza, F.A., Souza, T.M.L., Bosch, I., Yozwiak, N.L., MacInnis, B.L., Sabeti, P.C., 2017. Zika virus evolution and spread in the Americas. Nature 546, 411–415.

Moratorio, G., Henningsson, R., Barbezange, C., Carrau, L., Bordería, A.V., Blanc, H., Beaucourt, S., Poirier, E.Z., Vallet, T., Boussier, J., Mounce, B.C., Fontes, M., Vignuzzi, M., 2017. Attenuation of RNA viruses by redirecting their evolution in sequence space. Nat Microbiol 2, 17088.

Morlan, J.D., Qu, K., Sinicropi, D.V., 2012. Selective depletion of rRNA enables whole transcriptome profiling of archival fixed tissue. PLoS One 7, e42882.

Moudy, R.M., Meola, M.A., Morin, L.-L.L., Ebel, G.D., Kramer, L.D., 2007. A newly emergent genotype of West Nile virus is transmitted earlier and more efficiently by Culex mosquitoes. Am. J. Trop. Med. Hyg. 77, 365–370.

Myers, T.W., Gelfand, D.H., 1991. Reverse transcription and DNA amplification by a Thermus thermophilus DNA polymerase. Biochemistry 30, 7661–7666.

Nunes, M.R.T., Contreras-Gutierrez, M.A., Guzman, H., Martins, L.C., Barbirato, M.F., Savit, C., Balta, V., Uribe, S., Vivero, R., Suaza, J.D., Oliveira, H., Nunes Neto, J.P., Carvalho, V.L., da Silva, S.P., Cardoso, J.F., de Oliveira, R.S., da Silva Lemos, P., Wood, T.G., Widen, S.G., Vasconcelos, P.F.C., Fish, D., Vasilakis, N., Tesh, R.B., 2017. Genetic characterization, molecular epidemiology, and phylogenetic relationships of insect-specific viruses in the taxon Negevirus. Virology 504, 152–167.

Otte, A., Sauter, M., Daxer, M.A., McHardy, A.C., Klingel, K., Gabriel, G., 2015. Adaptive Mutations That Occurred during Circulation in Humans of H1N1 Influenza Virus in the 2009 Pandemic Enhance Virulence in Mice. J. Virol. 89, 7329–7337.

Quast, C., Pruesse, E., Yilmaz, P., Gerken, J., Schweer, T., Yarza, P., Peplies, J., Glöckner, F.O., 2013. The SILVA ribosomal RNA gene database project: improved data processing and web-based tools. Nucleic Acids Res. 41, D590–6.

Reidenbach, K.R., Cook, S., Bertone, M.A., Harbach, R.E., Wiegmann, B.M., Besansky, N.J., 2009. Phylogenetic analysis and temporal diversification of mosquitoes (Diptera: Culicidae) based on nuclear genes and morphology. BMC Evol. Biol. 9, 298.

Shi, M., Lin, X.-D., Tian, J.-H., Chen, L.-J., Chen, X., Li, C.-X., Qin, X.-C., Li, J., Cao, J.-P., Eden, J.-S., Buchmann, J., Wang, W., Xu, J., Holmes, E.C., Zhang, Y.-Z., 2016. Redefining the invertebrate RNA virosphere. Nature. https://doi.org/10.1038/nature20167

Shi, P.-Y., Tilgner, M., Lo, M.K., Kent, K.A., Bernard, K.A., 2002. Infectious cDNA clone of the epidemic west nile virus from New York City. J. Virol. 76, 5847–5856.

Tsetsarkin, K.A., Vanlandingham, D.L., McGee, C.E., Higgs, S., 2007. A single mutation in chikungunya virus affects vector specificity and epidemic potential. PLoS Pathog. 3, e201.

WHO, 2018. 2018 annual review of the Blueprint list of priority diseases [WWW Document]. who.int. URL http://www.who.int/emergencies/diseases/2018prioritization-report.pdf

Wilm, A., Aw, P.P.K., Bertrand, D., Yeo, G.H.T., Ong, S.H., Wong, C.H., Khor, C.C., Petric, R., Hibberd, M.L., Nagarajan, N., 2012. LoFreq: a sequence-quality aware, ultra-sensitive variant caller for uncovering cell-population heterogeneity from high-throughput sequencing datasets. Nucleic Acids Res. 40, 11189–11201.

Yamao, T., Eshita, Y., Kihara, Y., Satho, T., Kuroda, M., Sekizuka, T., Nishimura, M., Sakai, K., Watanabe, S., Akashi, H., Rongsriyam, Y., Komalamisra, N., Srisawat, R., Miyata, T., Sakata, A., Hosokawa, M., Nakashima, M., Kashige, N., Miake, F., Fukushi, S., Nakauchi, M., Saijo, M., Kurane, I., Morikawa, S., Mizutani, T., 2009. Novel virus discovery in field-collected mosquito larvae using an improved system for rapid determination of viral RNA sequences (RDV ver4.0). Arch. Virol. 154, 153–158.

Zakrzewski, M., Rašić, G., Darbro, J., Krause, L., Poo, Y.S., Filipović, I., Parry, R., Asgari, S., Devine, G., Suhrbier, A., 2018. Mapping the virome in wild-caught Aedes aegypti from Cairns and Bangkok. Sci. Rep. 8, 4690.

Zhang, X., Huang, S., Jin, T., Lin, P., Huang, Y., Wu, C., Peng, B., Wei, L., Chu, H., Wang, M., Jia, Z., Zhang, S., Xie, J., Cheng, J., Wan, C., Zhang, R., 2018. Discovery and high prevalence of Phasi Charoen-like virus in field-captured Aedes aegypti in South China. Virology 523, 35–40.

Zimmermann, L., Stephens, A., Nam, S.-Z., Rau, D., Kübler, J., Lozajic, M., Gabler, F., Söding, J., Lupas, A.N., Alva, V., 2018. A Completely Reimplemented MPI Bioinformatics Toolkit with a New HHpred Server at its Core. J. Mol. Biol. 430, 2237–2243.

